# STGram: Non-Invasive Visualization and Analysis of Circadian Rhythms Through Surface Temperature Monitoring

**DOI:** 10.1101/2024.05.21.595254

**Authors:** Shoi Shi, Tohru Natsume

## Abstract

Circadian rhythms, integral to physiological and behavioral processes, are influenced by environmental cues and developmental stages. This study explores the visualization and analysis of circadian rhythms through non-invasive monitoring of surface body temperature (STGram: Surface Thermo Deviations gram), focusing on the effects of jet lag in international travelers and the developmental progression of circadian rhythms in infants. Using a compact, wearable thermometric device, we collected data from adults experiencing jet lag and a 3-month-old infant over five months. Our analysis identified clear circadian shifts in travelers and illustrated the gradual establishment of circadian rhythms in the infant. These findings underscore the effectiveness of surface body temperature as a marker for circadian rhythm analysis, offering a valuable tool for understanding circadian dynamics and their impact on health. This methodological approach has significant implications for circadian rhythm research, health management, and the study of physiological development.

## Main

Organisms exhibit periodicity across various timescales, among which the circadian rhythm is notably conserved (1). The discovery of a molecular clock with a near-24-hour cycle underpinning behavioral rhythms marked a pivotal advancement in our understanding of temporal physiology (2,3). Subsequent research has elucidated numerous physiological processes influenced by the circadian rhythm, both upstream—such as the effects of light exposure and feeding—and downstream, including the sleep-wake cycle, dietary patterns, and thermoregulation (4–7). Consequently, circadian rhythms emerge as a fundamental regulatory mechanism for an array of physiological functions. It is now well-established that disruptions in circadian alignment with the external environment are implicated in a spectrum of diseases, extending beyond the traditionally recognized obesity and hypertension to encompass mental health disorders like depression (6,8–10). In response, interventions aimed at realigning circadian rhythms through cognitive-behavioral approaches have shown promise.

The quantification of circadian rhythms, traditionally reliant on core body temperature measurements or blood assays, offers precise snapshots but falls short in providing continuous, long-term insights (11,12). Recent advancements have seen a pivot towards leveraging wearable technology, such as smartwatches, for circadian rhythm visualization (13). These devices employ algorithms to analyze activity levels and sleep patterns, facilitating extended monitoring periods (14,15). Nonetheless, challenges persist, including the wearables’ size, battery life limitations, and the accuracy of rhythm detection via motion-based algorithms.

This study introduces an innovative approach by focusing on surface body temperature to predict circadian rhythms. We developed an algorithm, STGram, capable of analyzing time-series data from a compact, wearable device capable of over a month’s continuous temperature monitoring. Our findings demonstrate the device’s efficacy in visualizing jet lag and circadian rhythm development in infants, highlighting its potential as a simple, yet precise, tool for circadian rhythm assessment.

### Visualization of Temperature Time-Series Data

In our analysis, we employed a compact thermometric device with dimensions of less than ten inches (Figure 1a). The Hal-Share portable thermometer was selected for its minimal size and capability to perform extended duration temperature measurements exceeding one month. The collected raw temperature data was modified by applying a 10-minute averaging filter to enhance data smoothness (Figure 2b). Subsequently, we visualized the data spanning a one-month period, including the identification of maximum and minimum temperature values (Figures 1c,d). This analysis revealed that the median temperature values remained stable, exhibiting negligible fluctuations over the observed period (Figure 1d). Furthermore, a heat map representation of the monthly data delineated a discernible pattern of elevated surface body temperatures during nocturnal hours, coinciding with the sleep phases of the subjects (Figure 1e). This pattern underscores the potential of surface temperature monitoring as a non-invasive marker for circadian rhythm and sleep cycle analysis.

**Figure 1:**
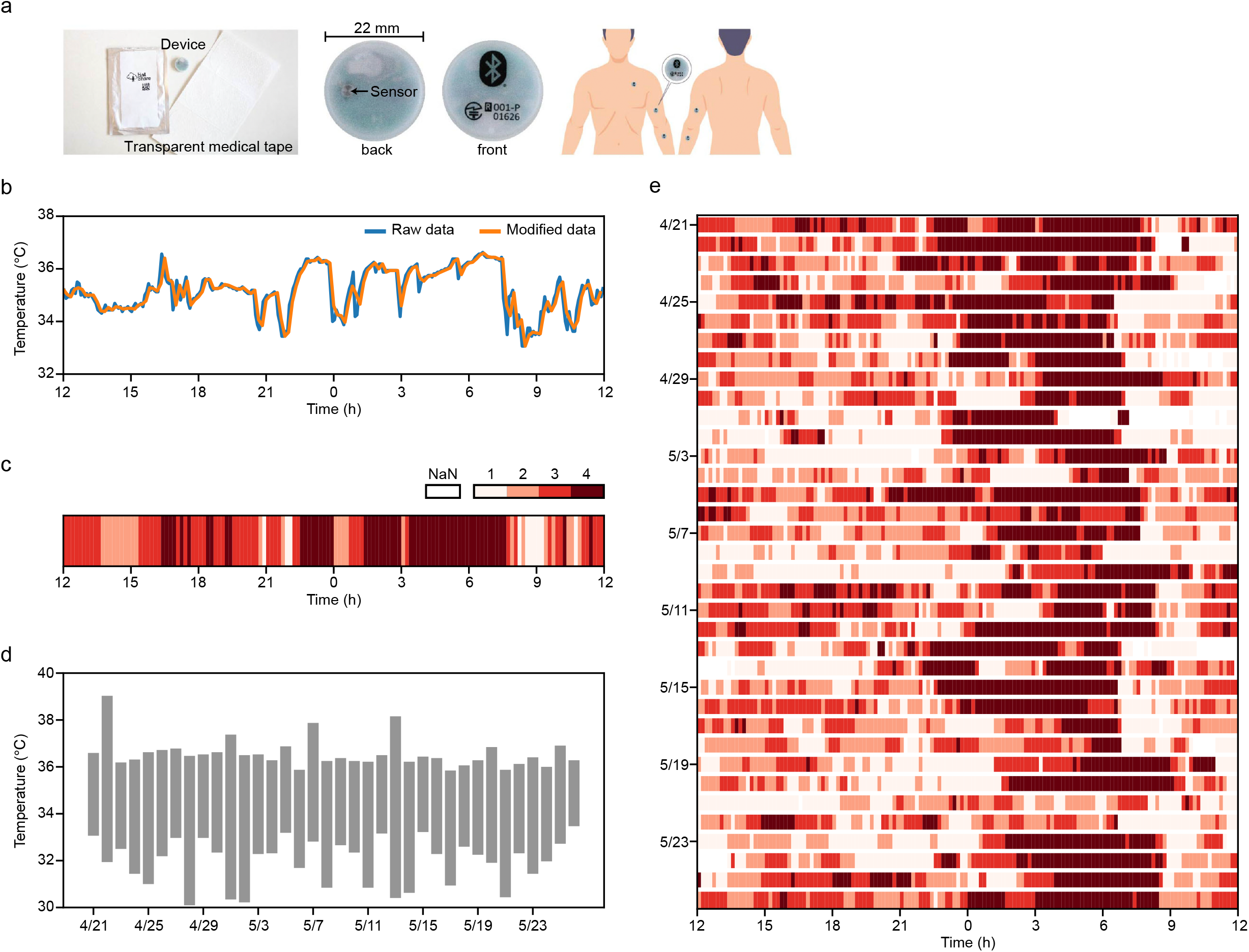
a. Diagram of the device. b. Time series of temperature data, showing both raw and modified data. The modified data are derived by averaging the raw data every 10 minutes. If no data are present within a 10-minute interval or if any data below 30°C are included, those data points are set to “nan.” These “nan” values are then replaced with the average of the non-”nan” data. c. Heat map of the temperature data for one day (n = 1), with darker red indicating higher temperatures. d. Plot showing the range of temperature data (n = 1), with maximum and minimum temperatures plotted for each day. e. Heat map representing approximately a month of data (n = 1).

**Figure 2:**
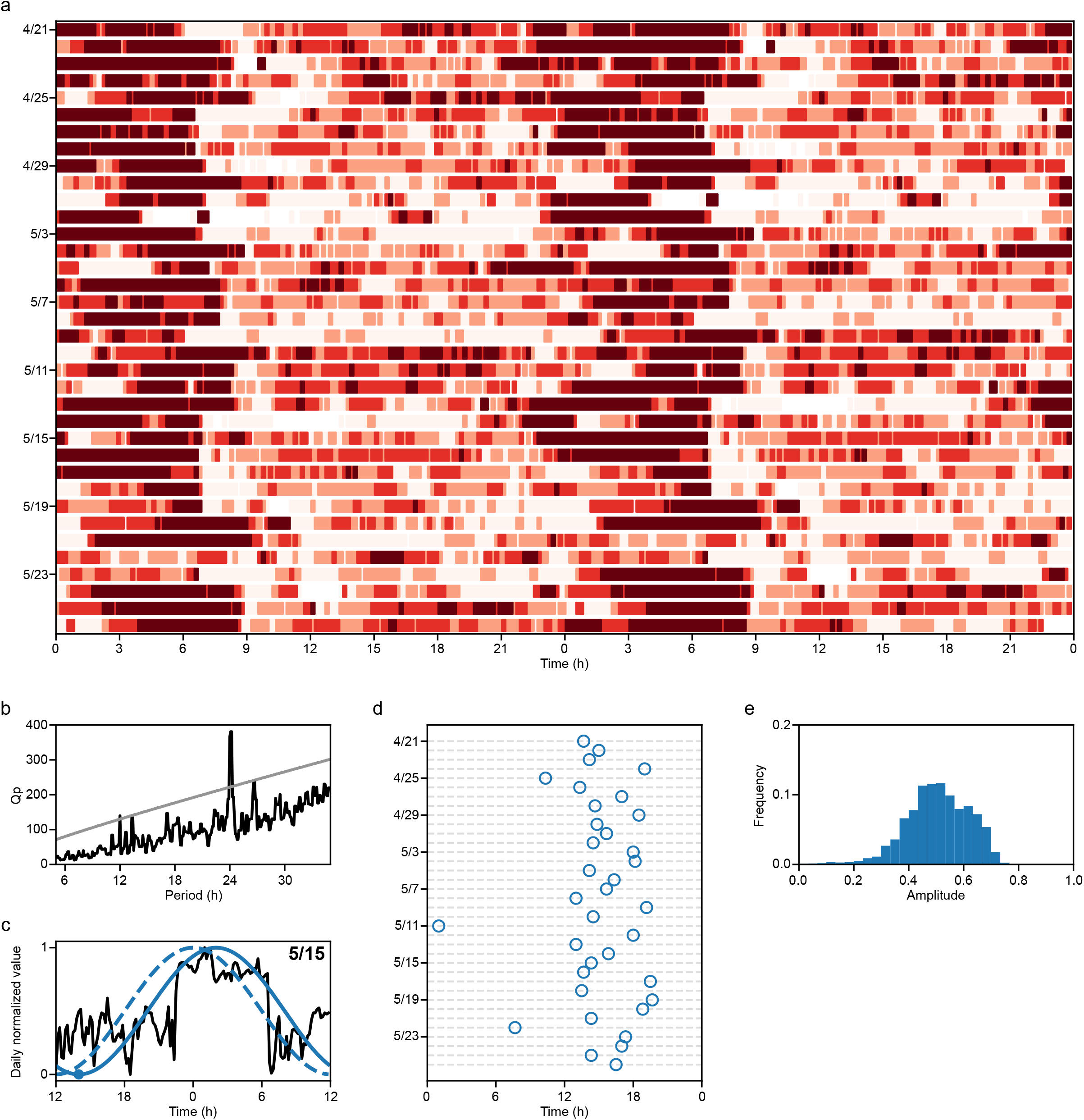
a. Double plot of heatmap (n = 1). b. Results of the chi-square periodogram are presented, where the black line represents the Qp values (a chi-square statistic), and the gray line indicates the 0.01 level of statistical significance, which ranges from 5.00 to 35.00 hours. c. This panel shows the body temperature time series data represented by a black line. The dashed and solid blue lines represent the van der Pol limit cycles. The dashed line corresponds to the cycle with its minimum point at noon, while the solid line is a curve fitted to the body temperature time series data, with a dot marking its minimum point. The phase is calculated as the duration between this minimum point and the preceding noon, which is 2.0 hours in this case. d. Phase plot for one month of data, providing a visual representation of the phase changes over the course of a month. e. Amplitude plot for one month of data, illustrating the variation in amplitude over the same month.

### Circadian Rhythm Analysis

To visualize circadian rhythms, we utilized a conventional double-plot representation (16) (Figure 2a). In instances of misaligned circadian rhythms, an anticipatory shift either to the right or left in the graphical representation would be expected; however, our analysis did not reveal such deviations. Furthermore, we developed an algorithm to quantify three critical metrics of circadian rhythm: period, phase, and amplitude (15,17) (Figures 2b-e). The period was determined by identifying the peak of the chi-square periodogram, a robust method for period detection outlined in the Materials and Methods section (Figure 2b). For phase calculation, we employed the van der Pol limit cycle model, a well-established approach in human circadian studies, to ascertain the circadian phase. The phase was defined as the interval from the minimum value on the model-fitted curve (illustrated as a blue point in Figure 2c) to the subsequent noon. Specifically, for the data in Figure 2c, the phase was 14 hours, which is a 2-hour difference compared to the control.

Given the inherent challenges in accurately measuring the true amplitude of circadian rhythms, particularly due to external influences such as light exposure, we opted for an alternative definition of amplitude. In this study, amplitude was quantified as the coefficient of variation (standard deviation divided by the mean) of hourly values (Figure 2e) (15). This measure was chosen to reflect the fluctuation intensity of a circadian-driven temperature output, offering insights into the intrinsic rhythmicity of body temperature regulation.

### Temperature Analysis Indicates Circadian Shifts Due to Jet Lag

Body temperature measurements were conducted on two international travelers utilizing the Hal-Share device over a one-month period. This interval encompassed stays in Japan (their original time zone) before and after traveling, as well as a week-long visit to destinations with time differences of -7 and -13 hours. The temperature data, visualized using double-plots, exhibited pronounced shifts corresponding to the new time zones, indicating hyperthermic phases (Figures 3a,b). The period analysis revealed a disruption or attenuation of the natural 24-hour cycle (Figure 3c), and the phase analysis of subject 2 demonstrated adjustments of -13 hours, aligning with the destination time zones (Figures 3d,e). These findings affirm the efficacy of employing surface body temperature data to compute and visualize external circadian rhythms, effectively mapping the impact of jet lag. Notably, for the subject 1, a gradual adaptation of the phase to the local time zone was observed (Figure 3a), underscoring the precision and quantitative capability of our algorithm in estimating circadian phase shifts based on surface body temperature. This adaptation highlights the potential of non-invasive temperature monitoring as a tool for assessing and understanding circadian dynamics in response to cross-time zone travel.

**Figure 3:**
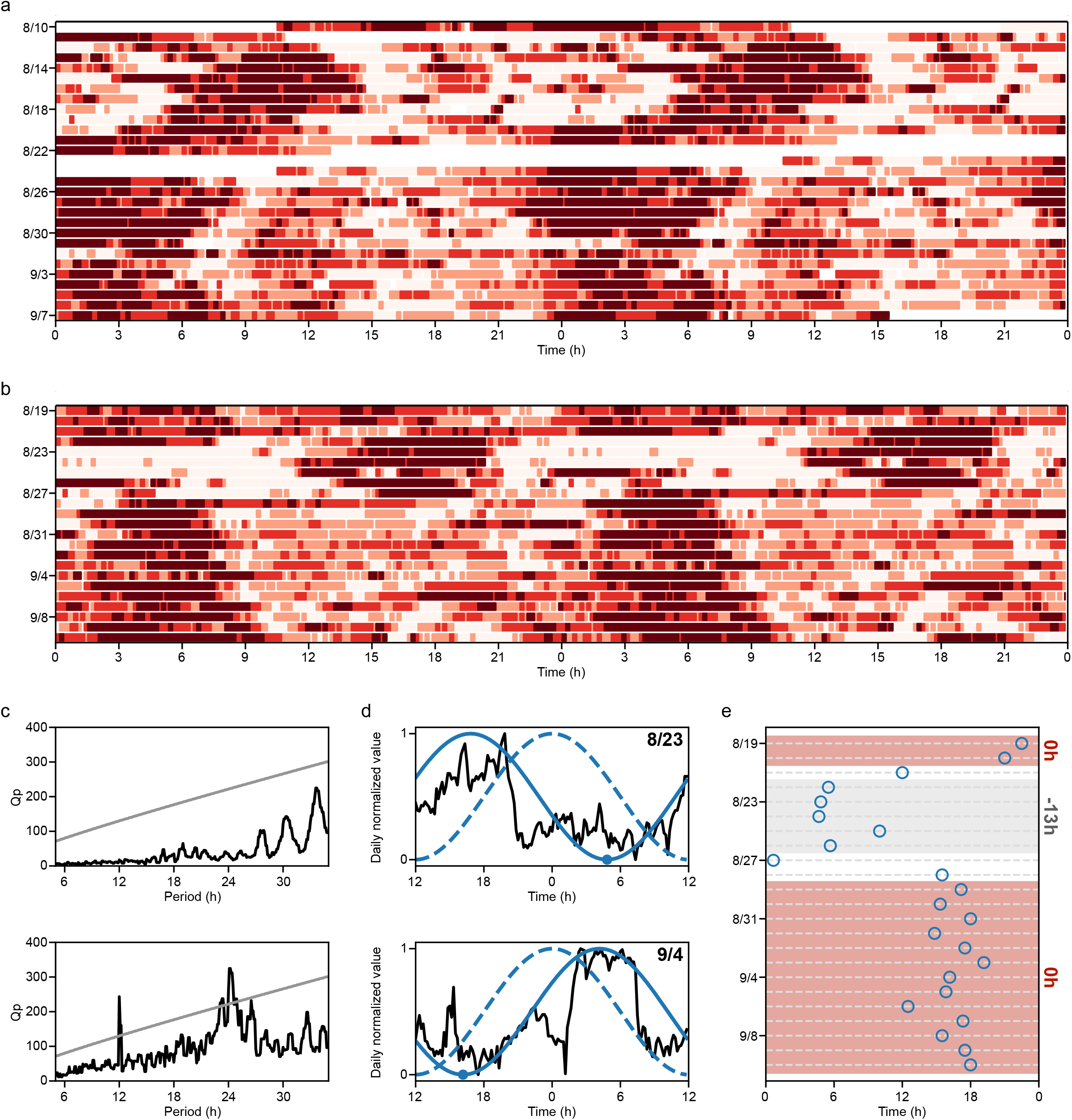
a. Double plot for subject 1. b. Double plot for subject 2. c. Period analyses for subjects 1 and 2. d. Phase analysis for subject 2 over two days, showing the timing of timezone changes and after returning to Japan. e. Phase transition, illustrating the phase moving from approximately 6:00 to about 18:00.

### Temperature Analysis Reveals Circadian Rhythm Development in Infants

Infants under the age of 3 to 6 months are known to possess underdeveloped circadian rhythms, which gradually synchronize with external environmental cues (18,19). By the age of 6 months, these rhythms typically become more established. In this context, our study aimed to determine whether our algorithm could effectively visualize this developmental process. We monitored an infant aged 3 months using the Hal-Share device across a five-month period. The analysis during the initial 3 to 6 months did not reveal significant 24-hour circadian cycles, and there was a noticeable variability in the phase measurements (Figures 4a-d, f-i, k-n). Conversely, by the age of 7 months, discernible 24-hour cycles were identified, and the phase variability markedly reduced (Figures 4e,j,o,p,q).

**Figure 4:**
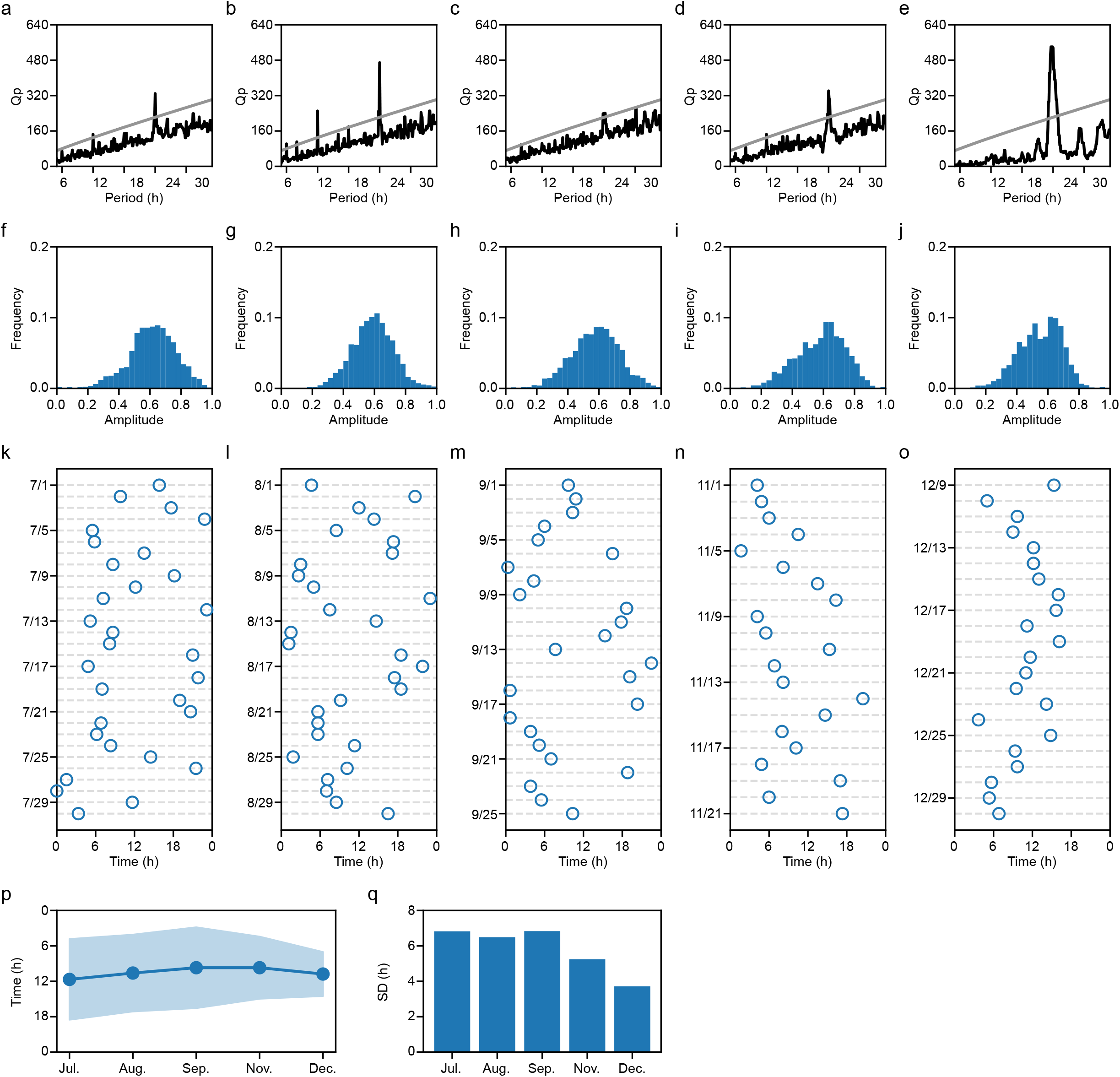
a-e. Period analysis for July, August, September, November, and December. All months show a peak around 24 hours, but peaks outside of 24 hours gradually diminish. f-j. Amplitude for July, August, September, November, and December. k-o. Phase trends for July, August, September, November, and December. p. Changes in the mean and variance of the phase. q. Plot of the standard deviation of p.

This pattern underscores the algorithm’s capability to capture the maturation of circadian rhythms from a phase of relative arrhythmicity to a more structured and predictable cycle. The findings highlight the developmental trajectory of circadian rhythms in infants, transitioning from an initial lack of a defined pattern to a more synchronized rhythm with the external day-night cycle.

## Discussion

This study presents a novel approach to visualizing and analyzing circadian rhythms through surface body temperature measurements using STGram. The findings underscore the utility of this method in capturing circadian dynamics across different scenarios, including the effects of jet lag on international travelers (Figure 3) and the development of circadian rhythms in infants (Figure 4).

Our analysis of international travelers wearing the small thermometer device revealed clear circadian shifts corresponding to changes in time zones, demonstrating the device’s effectiveness in visualizing the physiological impacts of jet lag. This capability to quantify adjustments in the circadian phase in response to cross-time zone travel offers potential applications in clinical and research settings, where understanding the extent of circadian disruption and its subsequent realignment is crucial (20,21). Moreover, the observed gradual adaptation of the circadian phase in one subject to the local time zone underscores the precision of our algorithm, highlighting its potential for personalized circadian rhythm management and adjustment strategies.

In the context of infant circadian rhythm development, our findings provide quantitative evidence of the maturation process of circadian rhythms from a nascent, arrhythmic state to a more defined and synchronized pattern (18,19). The absence of significant 24-hour cycles in infants aged 3 to 6 months and the emergence of these cycles by 7 months of age align with existing knowledge about circadian rhythm development. These results not only validate the capability of our algorithm in capturing these developmental changes but also suggest a non-invasive, practical tool for monitoring circadian health from an early age.

The study’s implications extend beyond these specific applications, highlighting the broader utility of non-invasive, continuous monitoring of circadian rhythms for health and disease management. For instance, the ability to track and analyze circadian rhythms could inform interventions for disorders where circadian misalignment is a contributing factor, such as sleep disorders, metabolic syndrome, and certain psychiatric conditions (22–24).

However, this study is not without limitations. The sample size, particularly in the infant development analysis, is small, and further research with a larger cohort is necessary to validate these findings. Additionally, the reliance on surface body temperature as a sole indicator of circadian phase may overlook other physiological or environmental factors influencing circadian rhythms. Future studies should consider integrating multiple physiological markers to enrich circadian rhythm analysis and provide a more comprehensive view of circadian health.

In conclusion, this research advances our understanding of circadian rhythms and their measurement, offering a novel tool for the non-invasive and continuous monitoring of circadian dynamics. The potential applications of this technology in health management, research, and personalized medicine are significant, paving the way for future explorations into circadian rhythm interventions and diagnostics.

## Methods

### Participants

The study recruited two distinct participant groups: international travelers and infants. International travelers were adults who planned cross-time zone travel of at least one week, with destinations having a minimum time difference of -7 and -13 hours from the original time zone. The infant group consisted of a single infant aged 3 months at the commencement of the study period, monitored over five months.

### Device and Data Collection

A compact, wearable thermometric device, specifically the Hal-Share portable thermometer (SUN・WISE Co., Ltd, Japan), was selected for temperature monitoring due to its small size (<10 inches) and extended data collection capability (over one month). Participants were instructed to wear the device continuously, allowing for the collection of surface body temperature data at regular intervals.

### Data Processing

Raw temperature data were collected and initially processed by smoothing, applying a 10-minute averaging filter to reduce noise. The processed data were then used for subsequent analysis, including the visualization of temperature variations and the determination of circadian metrics such as period, amplitude, and phase.

### Circadian Rhythm Analysis

Circadian rhythm analysis was performed using a custom algorithm designed to calculate three primary indicators: period, amplitude, and phase (also see previous study (15)). The period was determined through the chi-square periodogram method, with phase calculations based on the van der Pol limit cycle model. Amplitude was quantified as the coefficient of variation of hourly temperature values. This approach facilitated the identification of circadian shifts and the development of circadian rhythms, particularly in response to jet lag and during infant development.

### Statistical Analysis

Statistical analyses were conducted to assess the significance of observed circadian patterns and shifts. This involved comparing circadian metrics before and after travel for the traveler group and across different ages for the infant.

## Data availability

Data will be made available upon reasonable request.

## Code availability

The source code will be made available upon reasonable request.

## Acknowledgements

We appreciate all the members of Shi Laboratory in IIIS and Yasushi Senda for providing the device. This work was supported by the Japan Society for the Promotion of Science (JSPS) Grants-in-Aid for Scientific Research (KAKENHI) (20H05894, 20H05903, 21K15136, 22K21351, 23H02518A, 23H02663, and 23K18147 to S.S.), Japan Science and Technology Agency (JST)-Mirai Program (JPMJMI22J5 to S.S.), the Chugai Foundation and the Japan Foundation for Applied Enzymology to S.S., and the Top Runners in Strategy of Transborder Advanced Researches (TRiSTAR) by the MEXT to S.S.

## Author contributions

Study design was done by SS and TN. Data analysis, manuscript writing, and figure preparation were carried out by SS.

